# Transcriptomic responses to endurance exercise training in rats

**DOI:** 10.1101/2025.07.11.664421

**Authors:** Conrad Oakes, Lior Pachter

## Abstract

The bio-molecular changes of exercise, and how to best optimize them for improved performance, are an important human health research question. A recent study by the Molecular Transducers of Physical Activity Consortium (MoTrPAC) used a cohort of Rattus norvegicus to produce a whole-organism molecular map of the temporal effects of endurance exercise training. This dataset, encompassing hundreds of assays across 19 different tissues, can be used to evaluate the predictive power of gene expression, understand isoform-level changes in response to exercise, and with modern tools can be examined for associations with viral infection. Our analysis of the RNA-seq data reveals that gene expression can accurately predict the amount of exercise a rat was trained in. Additionally, we find biologically relevant isoform-level differences in expression that are masked in gene-level analysis. Finally, we find a potential novel virus that may negatively impact physiological measurements. This more comprehensive analysis provides a blueprint for directing similar analyses with respect to physiological perturbations across organisms.

## Introduction

Exercise is known to confer a variety of health benefits,(1–3) however, the precise molecular effects of exercise and how they translate into physiological changes are poorly understood (4). Key questions include how to best optimize diet, amount of exercise, the type of exercise, and sleep when undergoing a training regime. Of particular importance is whether molecular measurements can provide a roadmap to “precision health,” despite high variance between individuals. The search for biomarkers that provide clues to better understand the interplay between exercise and molecular biology could also have a significant impact on the prevention and treatment of injuries related to exercise, an important issue for athletes.(5, 6)

A recent study by the Molecular Transducers of Physical Activity Consortium (MoTrPAC) investigated the molecular effects of exercise in Rattus norvegicus,(7) and identified thousands of differentially expressed genes across tissues, as well as associations between exercise and other genomic modalities. The gene expression analysis was based on assaying 850 samples covering 17 tissues in 50 inbred Fischer 344 rats, along with an additional 25 samples covering each of the gonads (equaling 50 more in total). Despite the publication of an extensive analysis, the authors (7) did not use the MoTrPAC data to assess the predictive power of gene expression for extent of exercise, nor was the data used for a high-resolution analysis of isoform associations with exercise.

In analyzing the data with modern tools, we have discovered that gene expression can be an effective predictor of exercise. In a comprehensive evaluation of the data, we also find that some rats may have been infected with viruses, confounding associations with exercise. Furthermore, in investigating isoform changes with exercise, we find that exercise may substantially alter post-transcriptional modifications and that these may play a crucial role in physiological differences.

In what follows, we present a complete analysis of the MoTrPAC data and provide open-source code and notebooks that provide a blueprint for conducting such analyses, including with respect to other physiological perturbations, in humans.

## Results

### The MoTrPAC genomics corpus

The MoTrPAC gene expression data consists of experiments across 18 tissues in 50 rats (Figure 1a). The experiments conducted include bulk RNA-seq via an assay that included labeling molecules with unique molecular identifiers (UMIs), bulk ATAC-seq, Methyl-seq, acetylproteomics, metabolomics and lipidomics, phosphoproteomics, global proteomics, ubiqutiylome, and multiplexed immunoassays,. The experiments were conducted in rats that had exercised for 1 week, 2 weeks, 4 weeks, and 8 weeks, as well as control rats that were sacrificed after 8 weeks of no exercise. Each time point was interrogated in 5 male and 5 female rats, though the same rats were not sampled for every -omic sample (Supplemental Figure 1). Additionally, for rats in the control, 4 week, and 8 week cohorts, the changes in VO_2_ max performance, percent body fat, percent lean mass, and body weight over the course of the experiment were calculated.

**Fig. 1.**
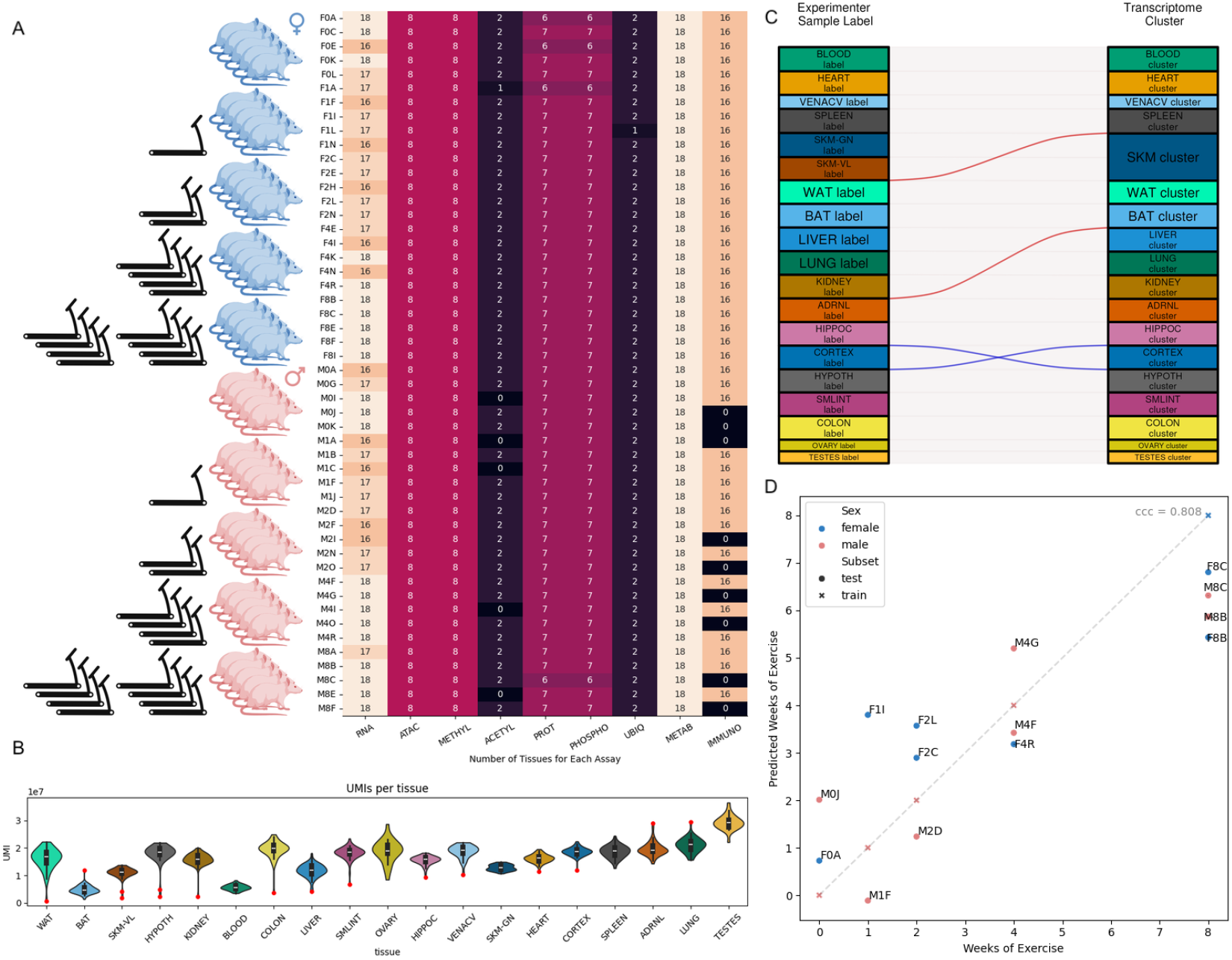
Linear regression results. A) An overview of individuals and their associated number of -omics assays post-QC. B) Violin plots of the UMIs per tissue from RNAseq results. Red dots are samples more than 3 standard deviations from the mean that were discarded from subsequent analyses. C) Alluvial plot of the difference between the original sample label and the identity of the Leiden cluster that the sample was assigned to. Red alluvia indicate samples that were incorrectly labeled and discarded; blue alluvia indicate two samples that were swapped. D) Linear regression results.

To examine the RNA-seq data, we processed it with kallisto-bustools (8), which are a state-of-the-art suite of tools for bulk and single-cell RNA-seq analysis that allows for counting unique molecules via UMIs (see Methods). A total of 21 samples were unavailable for reprocessing due to being discarded by the original study (all week 1 and 2 Vena Cava samples due to possible contamination and a single ovary sample for unexplained reasons), leaving the full dataset as 879 different bulk RNA samples.

We discarded an additional 14 samples whose UMI counts deviated by more than 3 standard deviations from the mean for that tissue type (Figure 1b); 10 of these had been discarded by the original analysis after being identified as outliers in a PCA analysis. We performed additional Leiden clustering of the samples and found that 4 samples were clustered in tissues different from their labels (Figure 1c). Three of these had been discarded as outliers in the original PCA analysis.(7) The sample that had not previously been identified as a sample to remove was a nominally hippocampus sample clustered in the hypothalamus cluster, while at the same time a nominally hypothalamus sample clustered into the hippocampus cluster. Because both samples were from the same individual rat, this appears to have been a clerical error resulting in swapped labels of the samples. We therefore relabeled the samples, thereby rescuing one, and included them in our downstream analyses.

### Prediction of exercise from molecular data

To directly address the question of whether gene expression can serve as a predictor of duration of exercise, we developed a generalized linear model regularized with ridge regression constraints. The model made use of the transcriptional profile of randomly selected subset of 36 rats across 9 different tissues. The input matrix for this model was created by, for each individual rat, flattening the matrix of gene counts for each tissue into a vector. In this way, an individual rat’s transcriptomic profile contains a different entry for a given gene for each tissue. This methodology was chosen with the idea that gene expression profiles may differ by tissue, so summing them is ineffective for getting an accurate overview of the expression landscape. Additionally, while samples were normalized for read depth, the log was not taken, as this would mean that linear regression is no longer calculating a true linear relationship between gene expression and time. For a given time point and sex, 2/3 of the samples were used for training and the remainder were used for testing. The result was a model that had a concordance correlation coefficient of.81 on the testing set (Figure 1d). Two of the top 10 most impactful gene coefficients of the overall linear regression were for the two types of skeletal muscle, and those tissues were also the most predictive when tissues were evaluated individually (Supplemental Figure 2).

Of the physiological measurements collected by the MoTrPAC, only the change in percent body fat could be effectively predicted by gene expression (Supplemental Figure 8), despite this being a rather noisy metric when viewing the change over time. The genes driving this predictive power appeared to make logical sense, with myoglobin expression in skeletal muscle predicting low body fat and more weeks of exercise, while Scd, a gene involved in fat differentiation, expression in brown adipose tissue predicting high body fat and fewer weeks of exercise.

When using the same method on the other assayed omics data, we found that proteomics was the best predictor. This analysis used the post-processing / QC results of the original study.(7) Using the same methodology to flatten the sample by feature matrix by individual, we analyzed all 9 different omics from the original study both at the same time and separately (Supplemental Figure 6). Proteomics and Acetylomics proved the most effective, which aligns with the original paper finding those to have some of the highest percent differentially expressed features. Methyl-seq and ATAC-seq had the worst predictive outcomes, again aligning with which -omics the original paper found had the fewest differential features proportionally. These trends were identical even when subsetting the features to those that corresponded to the most predictive gnes according to the RNA-seq regression (Supplemental Figure 10). However, this is not indicative of a lack of variability in the dataset, as both the Methyl-seq and ATAC-seq datasets were differential by tissue (Supplemental Figure 7).

We also assessed the predictive power of gene expression for exercise duration using the scVI (9) latent space representation (see Methods). We found that the latent space scrambles the information associated with exercise, leading to a loss of predictive power (Supplemental Figure 5). This was also the case when using the built-in batch correction methods to remove the effects of tissue/sex; in fact, on the 2D PCA plots the samples were still clustered by tissue (Supplementary Figure 4). This result is consistent with results of Antonsson et al.(10) which investigated the use of scVI for batch correction, and found that variational autoencoders, while possibly useful for data compression, may inadvertently remove significant amounts of biological signal.

### Isoform-level responses due to exercise

In addition to regression, we performed differential gene analysis using pyDeseq2(11) to find differential genes and isoforms within the dataset. The DESeq2 model(12) accounted for tissue, time, sex, and the interaction terms of tissue over time and sex over time. Using gastrocnemius skeletal muscle as a case study, we found that our model not only identified new differential genes over time but also recapitulated some of the results of previous work (Figure 2a-b).

**Fig. 2.**
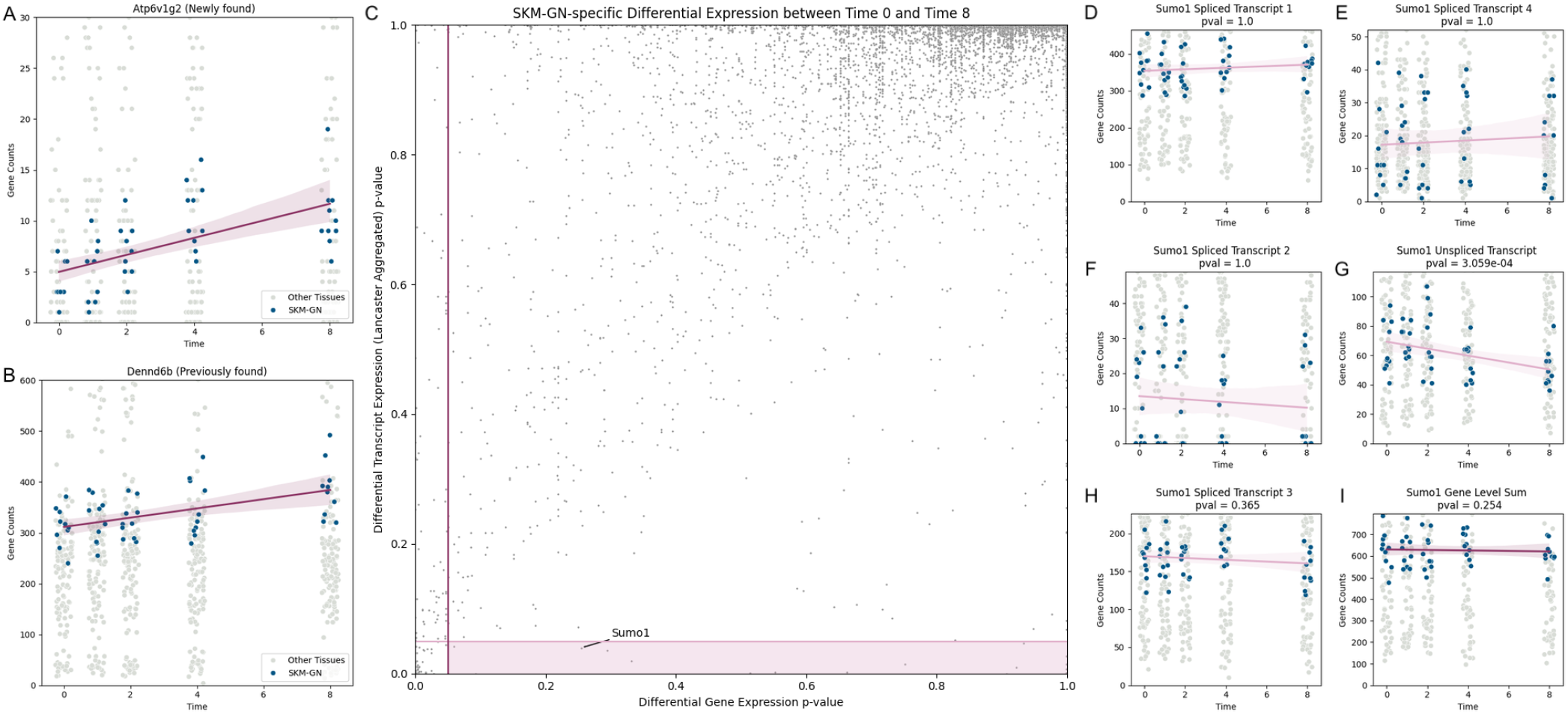
Isoform results A) Expression over time of gene X, previously found to be differential B) Expression over time of gene X, previously found to be differential C) Lancaster aggregated p-value versus Gene level p-value for the comparison of the interaction term of SKM-GN and Time D-I) Expression over time of the various transcripts of gene SUMO1

Next, we generated a similar model using the various transcript isoforms, rather than only the gene-level sums. To compare how this differed from the gene-level, the p-values for the transcripts of each gene were Lancaster aggregated using a weighting based off the log of gene counts for that transcript (13). We found 18 genes that were significantly differential at the isoform level but not at the gene level, and thus would have been missed with a purely gene level analysis, which had found 125 differentially expressed genes (Figure 2c).

An example of a gene that was transcript differential but not gene-level differential is SUMO, known to be a player in human response to resistance exercise (14). At the gene-level we found no noticeable changes in expression over time, whereas at the isoform level one of the transcript isoforms shows differential expression over time (Figure 2 d-g). Because the annotation of this exact rat gene is due to sequence similarity rather than direct investigation, it is possible that this may be indicative of a gene family-level change rather than an exact isoform change, though regardless it is still a change in expression missed by a standard gene-level analysis.

### Viral infection

We applied a recently developed method for identifying RNA virus sequences in RNA-seq data(15) via translated alignments to the PalmDB database.(16) A positive correlation (Pearson’s r of.49) was found between total virus counts in a sample and amount of time before sample collection (Figure 3a). Importantly, because the control rats were kept alive for the full 8 week time course, we included them with the 8 weeks of exercise rats for the purposes of this analysis. We then performed a differential expression analysis between week 8 and week 0 rats using pyDeseq2 in order to control for this apparent increase in virus counts over time. One virus-like sequence was found to be differentially expressed between the samples (Figure 3b). This virus-like sequence, u234187, had a statistically significant correlation with one of the genes reported as differentially expressed between time 0 and time 8 by the original authors (7) (Figure 3c). This gene, Tigit, is an immune expression gene.(17) When the raw reads corresponding to u234187 were run through BLAST,(18) the results aligned with several viral genomes (Supplemental Figure 12), namely ORF and SARS. Additionally, the rat that appears to have been infected by this putative virus, F0K, had the largest negative change in VO_2_ max among the time 0 rats (Figure 3e). Taken together, this implies that u234187 is a likely virus, and that the rat’s poor VO_2_ max was potentially due to viral infection rather than a sedentary lifestyle. Moreover, the likely false positive identification of Tigit as a gene differential due to exercise, highlights the importance of assessing the health of animals in studies attempting to link genomics or genetics data to physiological perturbations.

**Fig. 3.**
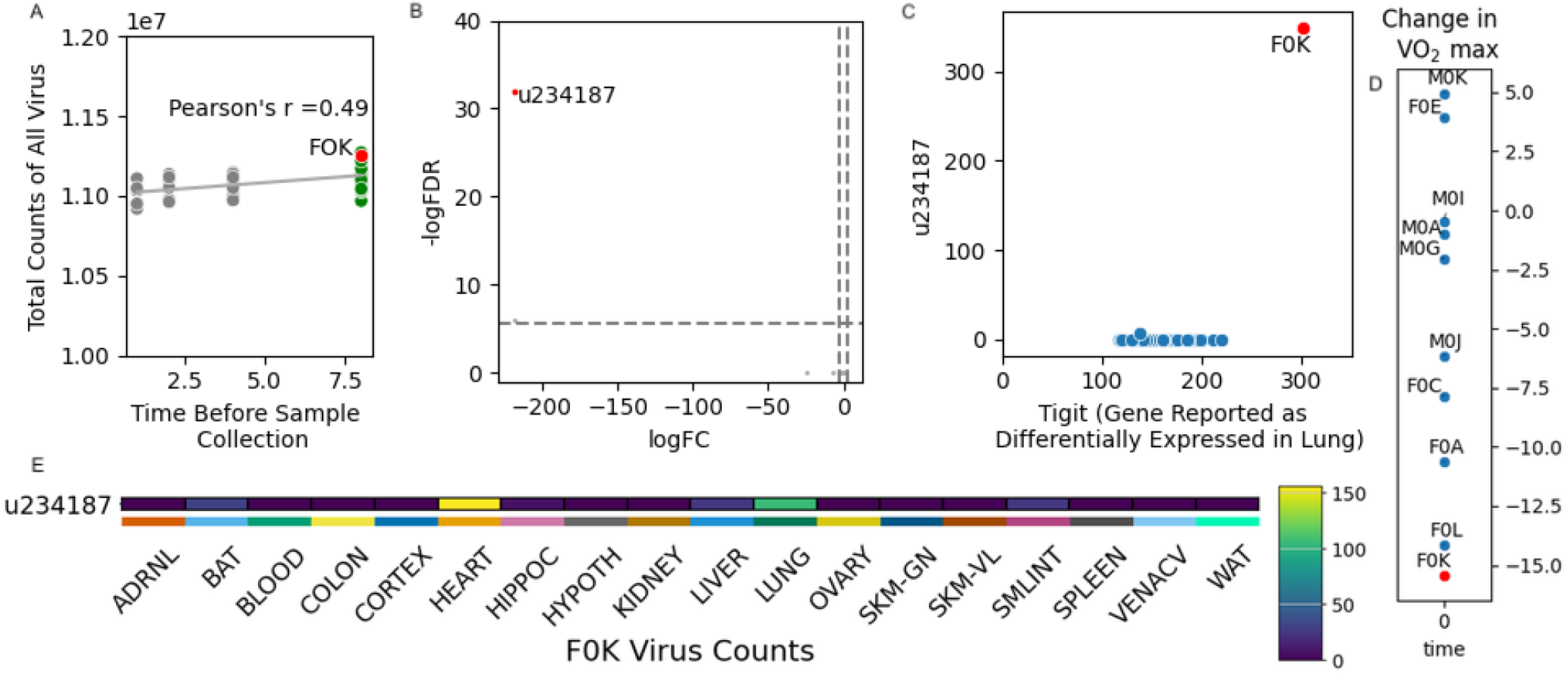
Virus sequence analysis results. A) Scatterplot of the time before sample collection and total virus counts in individual rats. B) Volcano plot of differential expression of virus counts between the 0 weeks of exercise and 8 weeks of exercise rats. C) Scatterplot between Tigit and virus u234187 expression. D) Change in VO_2_ max for all rats in the 0 weeks of exercise group. E) Heatmap of u234187 expression in rat F0K.

## Discussion

The explosion of sequencing data has resulted in a treasure trove of informative, yet under-analyzed data. The whole-organism RNA-sequencing done by the MoTrPAC is one such example of a large dataset with great potential for discovery. For example, thanks to the incorporation of unique molecular identifiers (UMIs) by the original researchers (7), the data can be mined for transcript-level changes due to exercise, an analysis not developed by the authors in their initial publication. We have also shown that comparisons between tissues and across the dataset are essential for comprehensive quality control. Importantly, the public release of data and metadata by the original researchers (7) made it possible for us to assess the predictive power of gene expression for the extent of exercise.

Specifically, we have shown that gene expression can accurately predict the amount of exercise rats undergo, and this result is robust to the machine learning approach used. In fact, we have shown that linear regression is effective for this task, whereas more sophisticated machine learning tools, such as variational autoencoders, may not be ideal for prediction. The fact that genes found to be strong predictors of exercise were also predictive when using their protein products but not their ATAC-seq data implies that the biomolecular changes associated with fitness are concentrated on post-transcriptional processes. In this case, the predictive power is perhaps more from RNA performing as a proxy for protein product rather than due to its regulation being the driving force.

While our attempts to use the ATAC-seq and Methyl-seq data for the same purpose resulted in no predictive power, it’s possible that a reanalysis of the raw data could improve their utility. Moreover, the ability to discern tissues by PCA in these datasets implies that there is some biologically informative variation in the data, and we believe the ATAC-seq data in particular may serve as a useful reference dataset for chromatin accessibility in rats.

We have also shown that changes in isoforms that could be biologically relevant are missed when looking at gene-level expression. This further emphasizes the necessity of using tools that can reach this level of resolution in order to better explain expression changes.

Finally, our exploration of putative virus expression found a potential novel virus that may have negatively impacted physiological measurements in the infected rat. Viruses are an underexplored aspect of RNA sequencing, and may constitute a potential confounding variable in many studies. As such, the fact that this virus-like sequence correlates strongly with Tigit, a previously found differential in time feature, and that the gene is expressed most highly in the specific infected rat, appears to be an example of a false positive result.

Future similar experiments could be further refined by, for example, more tightly controlling and tracking the amount of food consumed. Additionally, given that the physiological measurement most effectively predicted by gene expression was percent lean mass, it would be useful to compare to the gene expression of a rat with a similar phenotype but different underlying conditions (for instance, starvation) to decouple which aspects of gene expression that are guiding this prediction are truly related to exercise.

## Data and Code Availability

All code to download data and generate the main and supplementary figures is available at https://github.com/pachterlab/CP_2025, executable in Google Colaboratory notebooks.

## Acknowledgements

We thank Laura Luebbert for help and advice on virus detection in the RNA-seq samples analyzed, and Barbara Wold for suggesting an examination of the ATAC-seq in genes predictive of fitness. We thank the Molecular Transducers of Physical Activity Consortium (MoTrPAC) for making their data freely available and clearly annotated. Cartoons used in Figure 1 were generated in BioRender.

## Methods

### Acquisition and preprocessing of RNA-seq data

Processed RNA-seq reads from (7) were obtained from the Sequence Read Archive. These fastq files containing the paired reads and the UMI barcode were then passed into the kallisto/bustools pipeline. Due to being a bulk RNA dataset with UMIs, a custom technology string had to be used in lieu of the standard kb count command. For details and scripts on the pipeline, see https://github.com/pachterlab/OP_2025/blob/main/commandline_data_generation_scripts/Download_and_Count_SRR_Data.sh After counting, the resulting count matrices were combined into one h5ad file for downstream processing. This count matrix was passed into R for read length normalization with the tximport package. For details and scripts on the read length normalization, see https://github.com/pachterlab/OP_2025/blob/main/R_scripts/RNA_to_adata.Rmd The resulting count matrix was then loaded into scanpy for basic filtering. Sample UMIs were summed, then outliers for each tissue (defined as greater than 3 standard deviations from the mean for each) were discarded. Leiden clustering was then performed. The 4 samples whose labeled tissue identity did not align with their cluster identity were inspected. Of these 4, 2 were determined to be potentially contaminated samples and removed from downstream analyses, while the 2 were determined to be a label swap and so were retained but corrected before proceeding. For details and scripts on the initial filtering, see https://github.com/pachterlab/OP_2025/blob/main/analysis_scripts/Initial_RNA_Analysis.ipynb. For details and scripts on the analysis of the corrected data, see https://github.com/pachterlab/OP_2025/blob/main/analysis_scripts/Label_Correction_Reanalysis.ipynb.

### Acquisition of other -omic data

Fully processed count matrices from (7) were obtained from the MoTrPAC github repository. All of these files were used without modification. For details and scripts on the data download, see https://github.com/pachterlab/OP_2025/blob/main/R_scripts/OMIC_Data_Download.Rmd and https://github.com/pachterlab/OP_2025/blob/main/R_scripts/Methyl_Data_Download.Rmd.

### Linear Regressions

A generalized linear models were constructed using the scikit sklearn package in python with the ridge regression setting. For linear regressions using RNA the normalized, but not log, gene counts were used after concatenating count matrices on the sample level. For details and scripts on linear regression of RNA to predict extent of exercise, see https://github.com/pachterlab/OP_2025/blob/main/analysis_scripts/Linear_Regression_RNA.ipynb. For details and scripts on linear regression of the scVI latent space to predict extent of exercise, see https://github.com/pachterlab/OP_2025/blob/main/analysis_scripts/Linear_Regression_scVI.ipynb. For details and scripts on linear regression of -omics to predict extent of exercise, see https://github.com/pachterlab/OP_2025/blob/main/analysis_scripts/Linear_Regression_OMIC.ipynb. For details and scripts on linear regression of RNA to predict other physiological measurements, see https://github.com/pachterlab/OP_2025/blob/main/analysis_scripts/Linear_Regression_Physiological.ipynb.

### Differential Expression Analysis

The pydeseq2 package was used to perform differential expression analysis. The input model formula of (*tissue* + *sex*)* *time* was used to generate the DESeq model. For details and scripts on the differential expression analysis, see https://github.com/pachterlab/OP_2025/blob/main/analysis_scripts/DEseq.ipynb.

### Generation of Virus count matrix

Processed RNA-seq reads from (7) were obtained from the Sequence Read Archive. These fastq files containing the paired reads and the UMI barcode were then passed into the kallisto/bustools pipeline. Due to being a bulk RNA dataset with UMIs, a custom technology string had to be used in lieu of the standard kb count command. The virus sequences index used as a reference was generated with host masking in order to prevent false positives. For details and scripts on virus count matrix generation, see https://github.com/pachterlab/OP_2025/blob/main/commandline_data_generation_scripts/Virus_Download_and_Count_SRR_Data.sh. After counting, the resulting count matrices were combined into one h5ad file for downstream processing.

### Virus Quality Control

Viruses that corresponded to known contaminates from laboratory reagents were annotated and then removed for further downstream analysis. For details and scripts on this processing, see https://github.com/pachterlab/OP_2025/blob/main/analysis_scripts/Virus_Filtering.ipynb.

### Virus Analysis

Total virus-like counts were summed across all tissues in a specific rat. This summed count was then correlated with time until death. To control for this correlation, only control and week 8 rats were used for DESeq2 analysis. For details and scripts on this correlation analysis, see https://github.com/pachterlab/OP_2025/blob/main/analysis_scripts/Virus_Correlation_Over_Time.ipynb.

This comparison found 2 significant virus-like sequences after p-value adjustment. These virus-like sequences were then correlated with total gene counts, with significance being calculated post-Bonferroni correction. For details and scripts on this comparison and correlation analysis, see https://github.com/pachterlab/OP_2025/blob/main/analysis_scripts/Virus_DEseq.ipynb.

The raw reads corresponding to virus-like sequence u234187 were extracted and then run through the BLAST database to detect similar sequences. For details and scripts on the BLAST analysis, seehttps://github.com/pachterlab/OP_2025/blob/main/analysis_scripts/Virus_BLAST.ipynb.

