## Supplemental Figures for "Transcriptomic responses to endurance exercise training in rats"

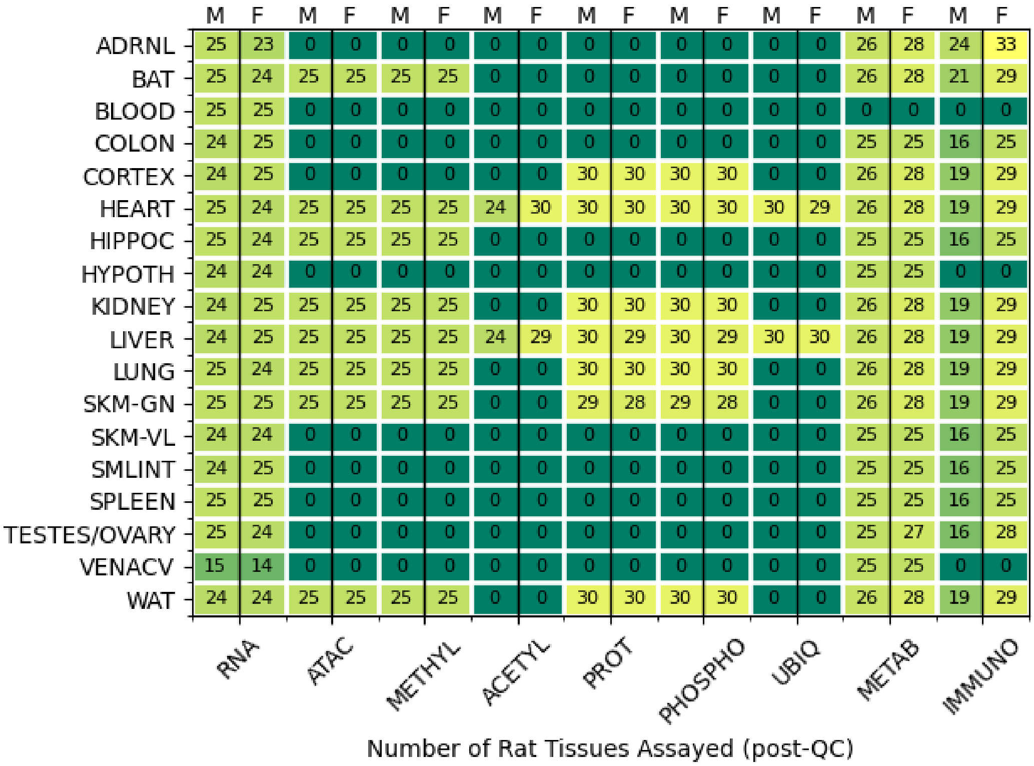


**Fig. 1.** Samples of Each Tissue Available for Each Assay post-QC. An overview of number of samples were collected and passed quality control for each assay in each tissue, split by sex.

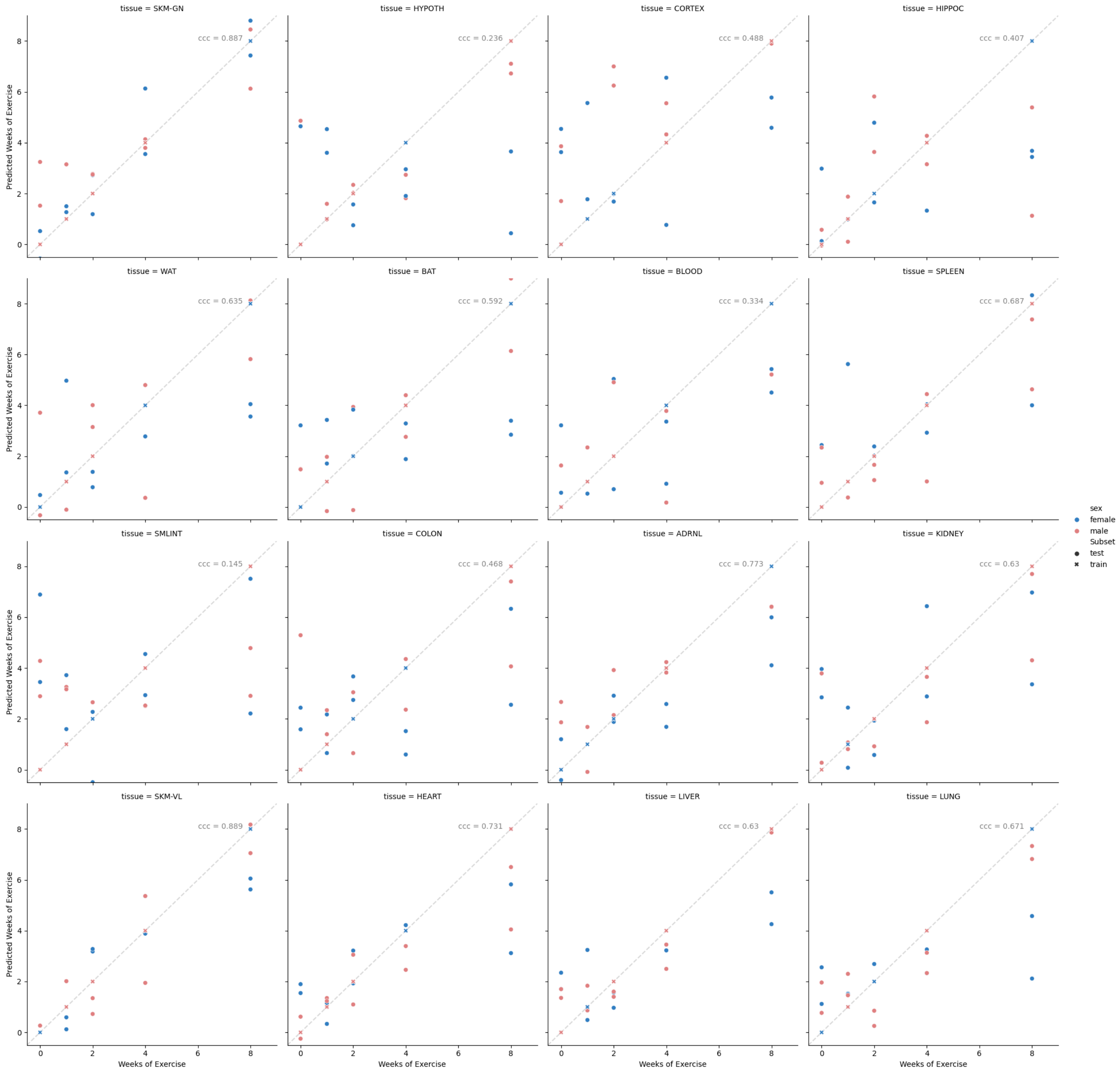


**Fig. 2.** Linear Regression Results for Individual Tissues. The results of generating a Generalized Linear Model to predict weeks of exercise using only the data from an individual tissue.

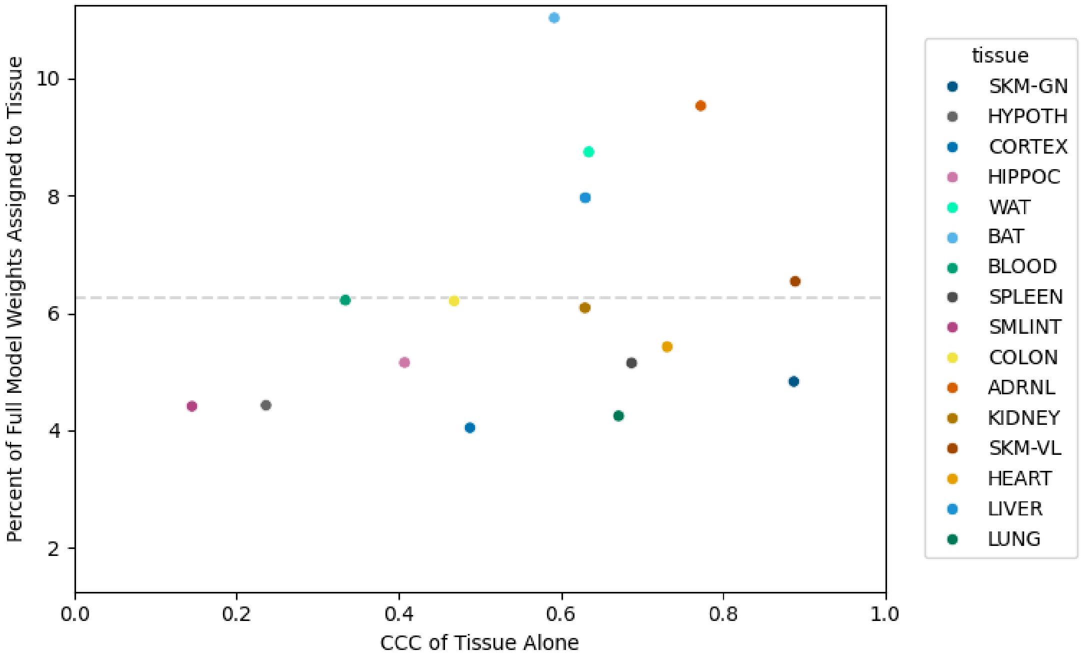


**Fig. 3.** Comparison of individual tissue Linear Regressions to overall, combined linear regression. The x-axis is the concordance correlation coefficient of each tissue when used to predict weeks of exercise with a GLM. The y-axis is the percentage of full model weights that were assigned to that tissue, with the dotted line indicating the value one would expect if each tissue contributed an equal amount of weight to the full model.

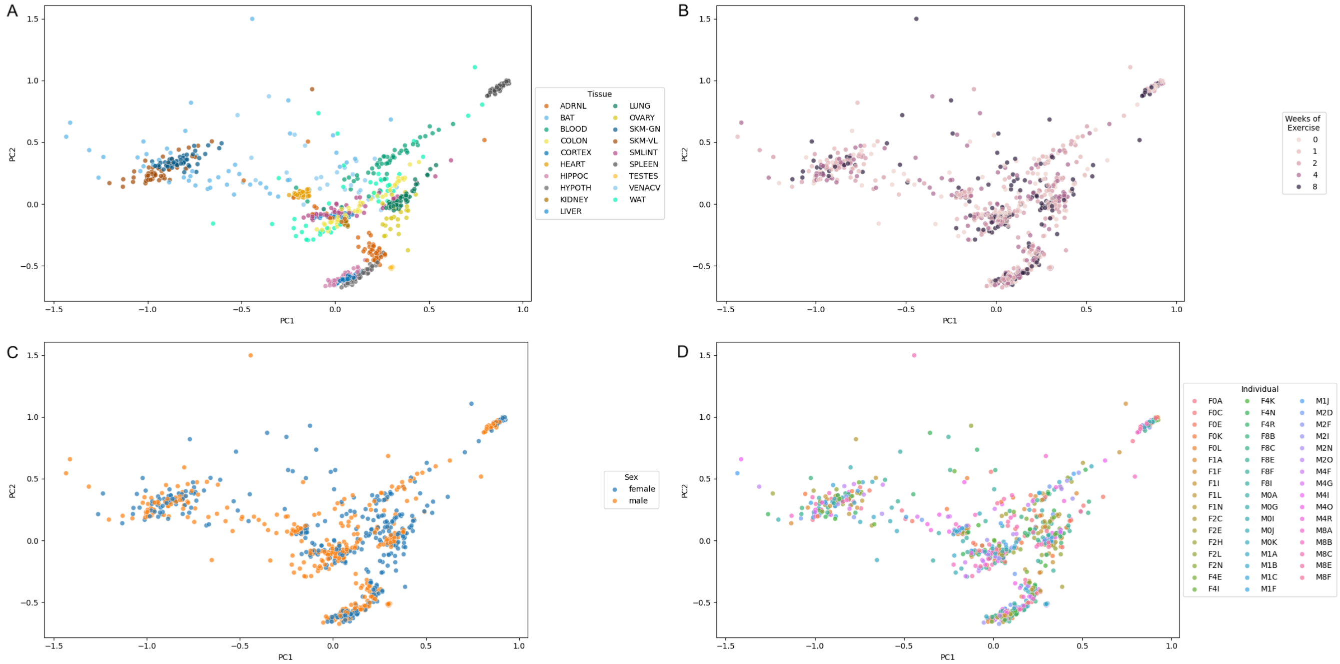


**Fig. 4.** PCAs of data post-scVI batch correction. PCA plots of the data post-scVI batch correction for tissue and sex differences. Plots highlighted by A) Tissue, B) Time, C) Sex, and D) Individual.

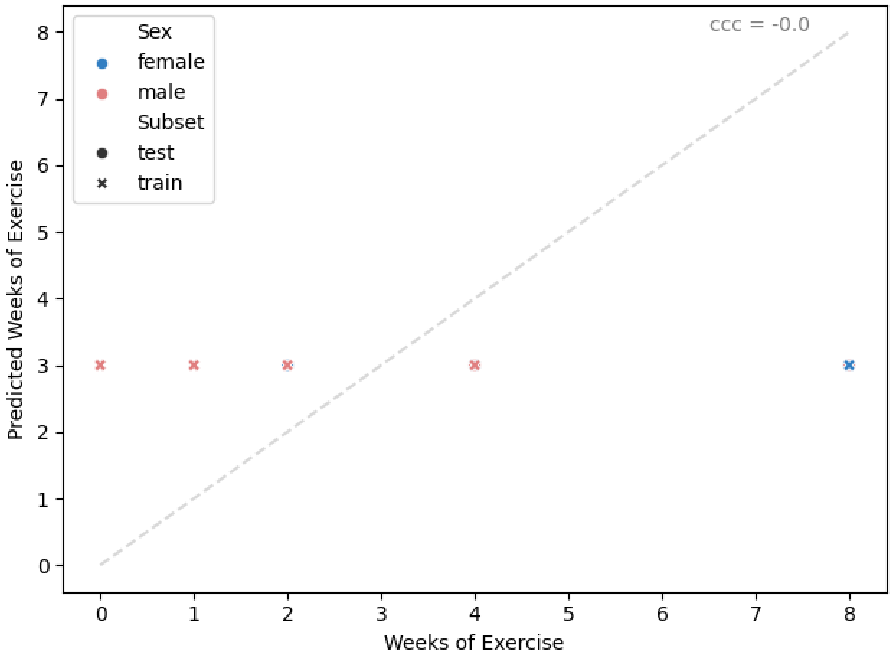


**Fig. 5.** Linear Regression to Predict Weeks of Exercise using scVI Latent Representation. The results of generating a Generalized Linear Model to predict weeks of exercise using the latent space representation of the data.

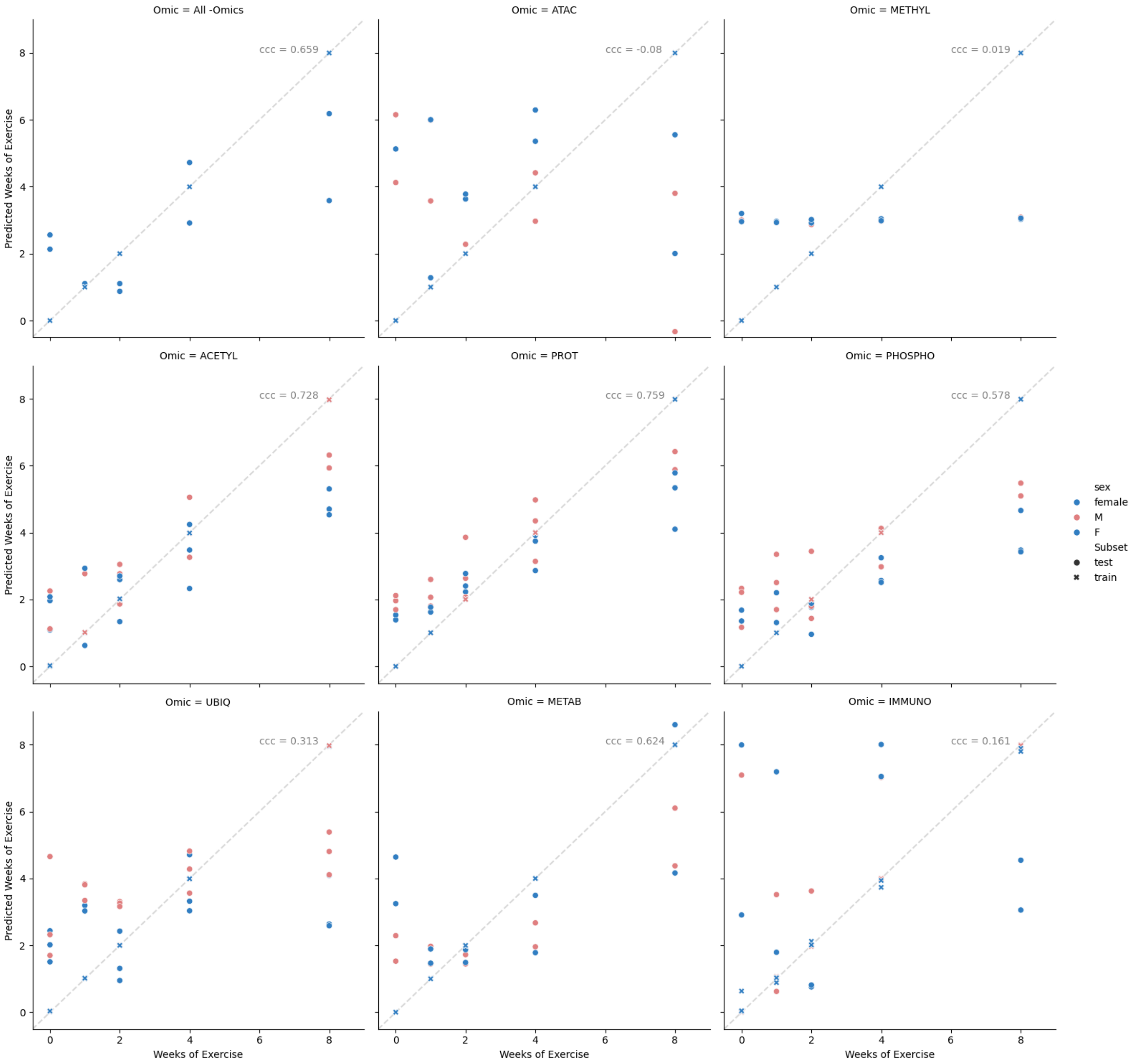


**Fig. 6.** Linear Regression to Predict Weeks of Exercise using non-RNAseq assays. The results of generating a Generalized Linear Model to predict weeks of exercise using all assay data together (limited to only female heart tissue) or each assay individually (using all tissues).

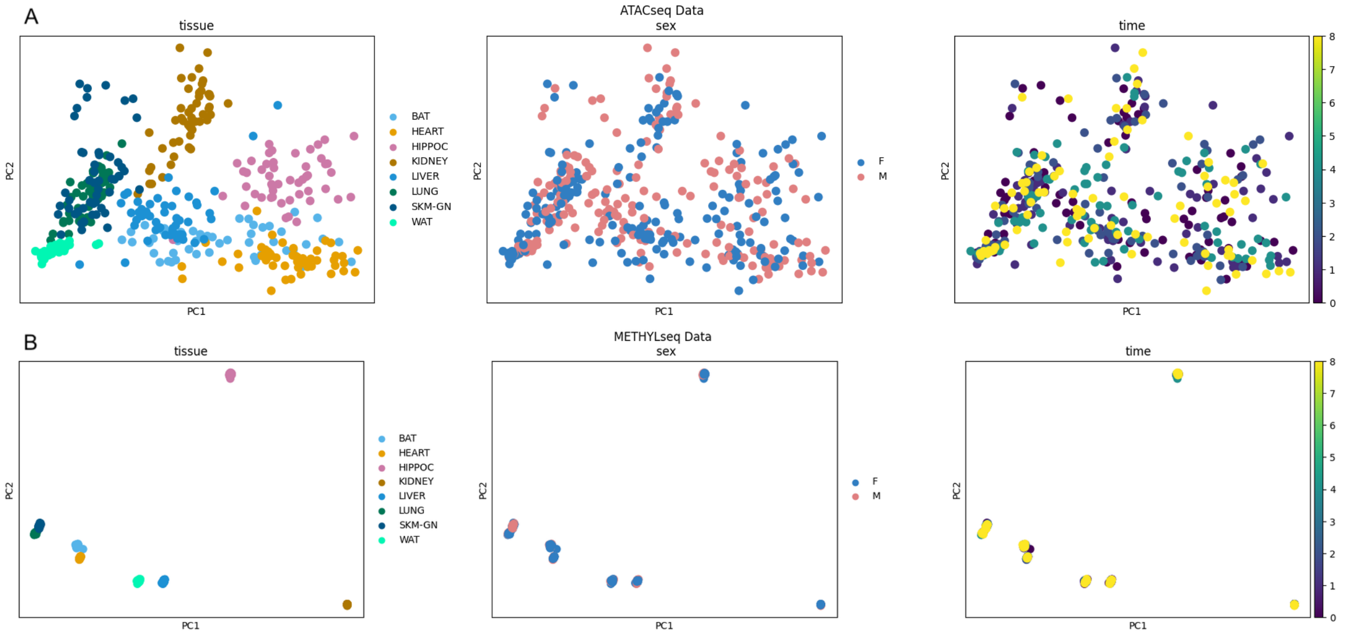


**Fig. 7.** PCA of Methyl-seq and ATAC-seq data. PCA plots of A) ATACseq data and B) Methyl-seq data, highlighted by tissue, sex, and time.

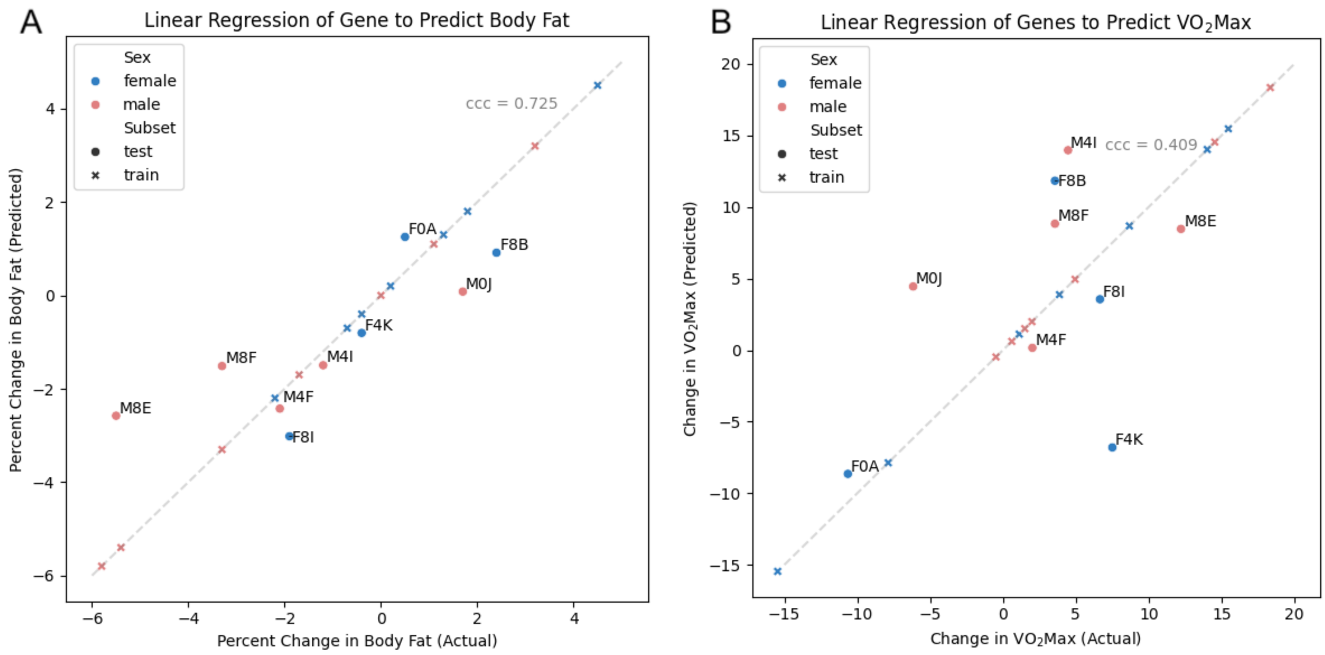


**Fig. 8.** Linear Regression to Predict Physiological Measurements. The results of generating a Generalized Linear Model to predict A) Percent Change in Body Fat and B) Change in $VO_2$ Max.

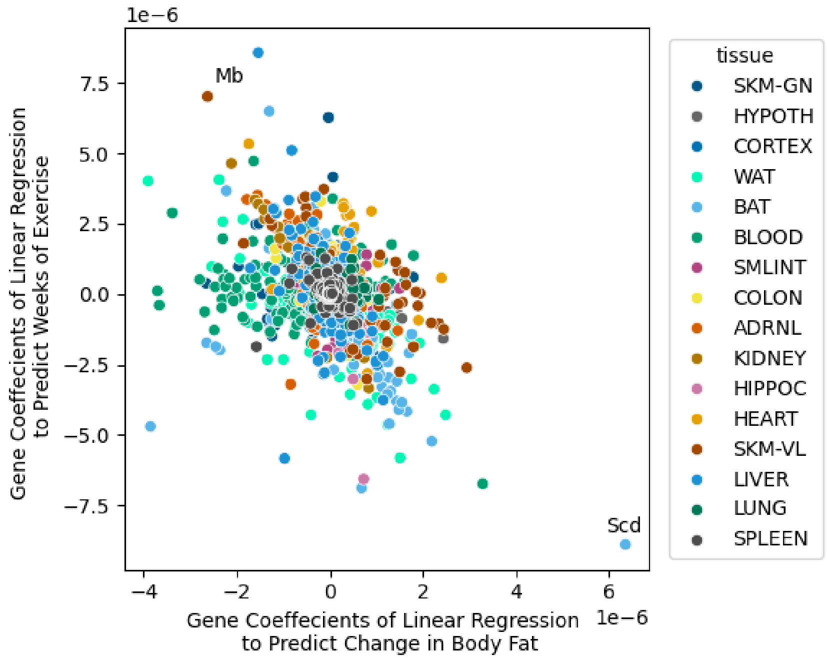


**Fig. 9.** Linear Regression Model Weights for Genes when predicting Percent Change in Body Fat vs when predicting Weeks of Exercise. The model weight coefficients for each gene when used to predict weeks of exercise (y-axis) or percent change in body fat (x-axis), colored by tissue.

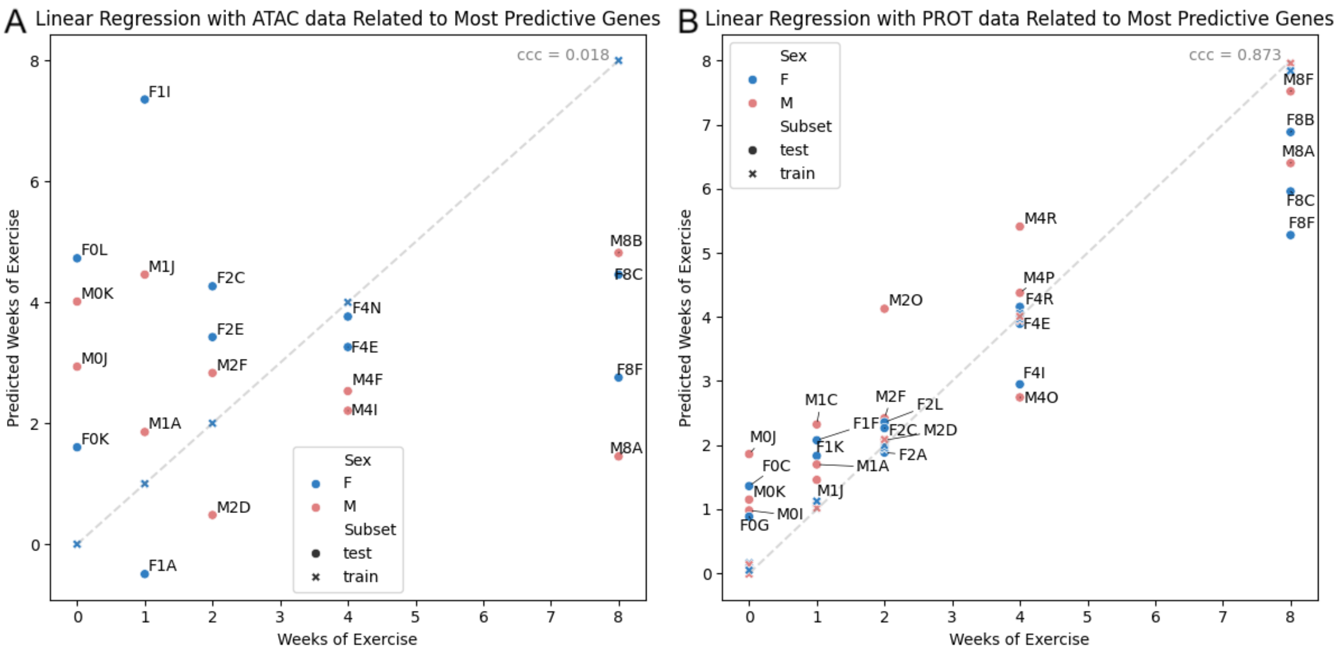


**Fig. 10.** Linear regression to predict weeks of exercise for A) ATAC data and B) PROT data after filtering for the genes most predictive in the RNA linear regression.

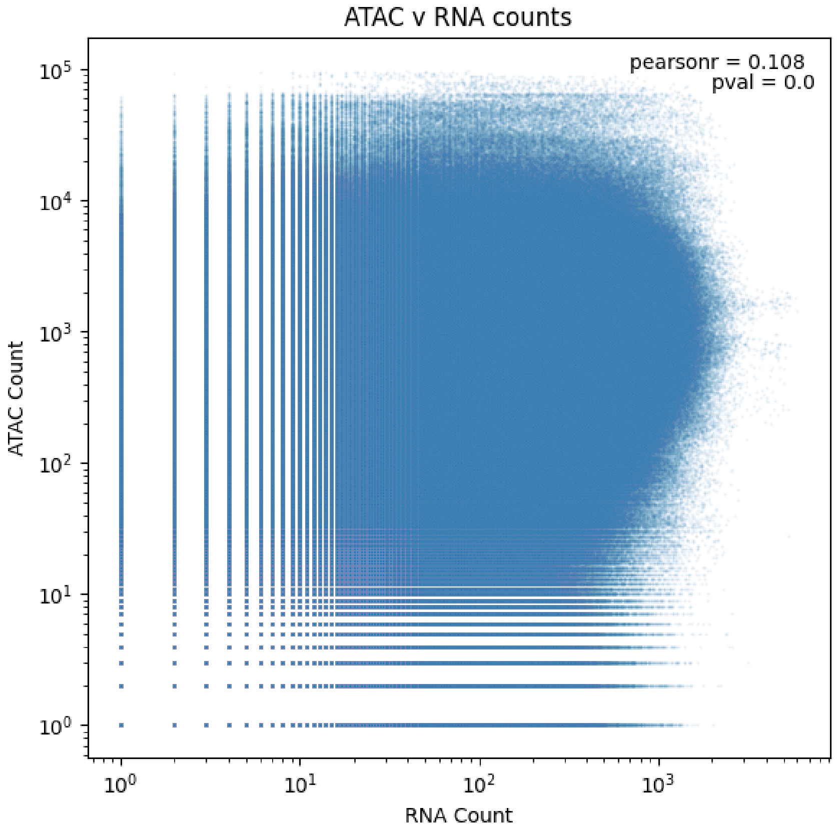


**Fig. 11.** For each gene, correlation between its RNA counts and ATAC counts for its related promoters within 1 kb.

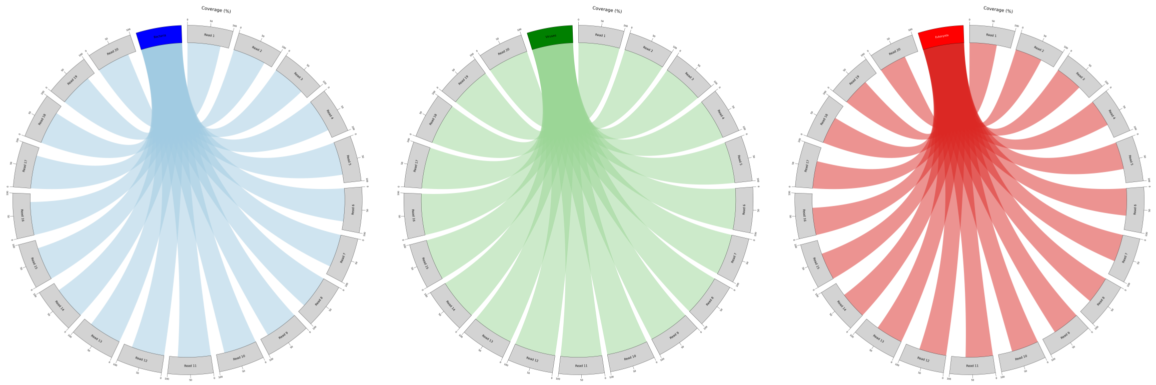

**Fig. 12.** pyCirclize (https://github.com/moshi4/pyCirclize) plots showing the BLAST results of sequencing reads for putative virus u234187 randomly selected from rat F0K. Each light grey sector corresponds to one sequencing read that links to the superkingdoms (red (eukaryotes), blue (bacteria), and green (viruses) sectors) based on its BLAST alignment results. The width of the connecting link indicates the BLAST alignment coverage percentage.
